# Loss of conserved noncoding elements likely shaped the evolution of regressed phenotypes in cavefish

**DOI:** 10.1101/2024.07.05.596787

**Authors:** Mohan Lal, Jui Bhattacharya, Kuljeet Singh Sandhu

## Abstract

The Mexican cavefish, Astyanax *mexicanus*, is a captivating model for probing cave adaptations, showcasing pronounced divergence in traits like vision, brain morphology, behaviour, pigmentation, and hypoxia tolerance compared to its surface-dwelling counterpart. Very few protein-coding variants are identified in cave-morphs, and the vast phenotypic gap between the two morphs remains inadequately explained. We investigated the noncoding genomes of teleosts and found that 3,343 conserved non-coding elements (CNEs) were independently lost in cave-morphs. These CNEs, confirmed in Zebrafish, displayed enhancer-associated histone modifications, possessed binding sites of neuronal transcription factors and interacted with cognate genes through chromatin loops. Genes crucial for eye and nervous system development were located adjacent to CNEs lost in cave morphs. Notably, these flanking genes were gradually downregulated during embryonic development of cave-morphs, contrasting with surface morphs. These insights underscore how compromised developmental pathways, stemming from the loss of distal regulatory elements, contribute to the regression of phenotypes in cave morphs.

**Article Summary:** Despite availability of genome sequences and allied datasets, the genetic underpinning of regressed traits of cavefish remains enigmatic. By aligning the genome sequences of teleosts, we identified thousands of noncoding elements specifically lost in cavefish, exhibited enhancer-associated hallmarks, and were enriched with the binding sites of neuronal transcription factors. Their cognate genes were associated with eye and nervous system development, and exhibited developmental downregulation in cavefish. This study highlights how the loss of regulatory elements impacted the cavefish evolution and adaptation.

## Introduction

Species of several phyla are adapted to cave environment and exhibit convergently diverged phenotypes like loss of vision, albinism and distinct neuroanatomy and behaviour, (Culver and Pipan 2019). The genetic basis of these regressed phenotypes is not entirely understood. Accumulating data of paired surface and cave morphs of teleost Astyanax *mexicanus* represent a unique opportunity to study the genomic basis of regressive evolution(Dowling et al. 2002). Vision, essential for surface habitat, is significantly regressed in cave morphs of Astyanax *mexicanus*. In cavefish embryos, the eye primordia are initially formed, but are subsequently arrested and progressively degenerated(Jeffery et al. 2003; Yamamoto et al. 2004). It has been shown that cavefish embryonic lens is destructed through the process of apoptosis(Jeffery and Martasian 1998; Alunni et al. 2007). But, this is not the only mechanism leading to vision loss in cavefish. Firstly, the optic vesicles in cavefish embryos, before any degeneration, are significantly smaller than that of surface morph(Jeffery et al. 2003). Secondly, the optic cup is absent in cavefish embryos(Jeffery et al. 2003). Therefore, it is tenable to hypothesize that the mechanisms other than apoptosis might independently contribute to regressed eyes in cavefish. The cave morphs also have smaller brain mass than surface morphs, specifically the optic tectum and periventricular grey zones(PGZ) are significantly regressed in cave morphs. Further, the cave morphs also have complex alteration in craniofacial structure, particularly the bone fragmentations and fusions through unusual ossification process(Powers et al. 2018). Several of these craniofacial changes have arose independently from the regression of eyes(Yamamoto et al. 2003). Some craniofacial changes, like protruded lower jaw, are likely constructive adaptations for altered feeding behaviour(Powers et al. 2023). Cave morphs exhibit significantly diverged behaviour from that of surface morphs. This includes loss of circadian rhythm, handedness in swimming, greater sensory perception, greater foraging, and social avoidance behaviour(Yoshizawa 2015). These behavioural changes can partly be attributed to the enlargement of hypothalamus and other neuroanatomical changes in cave moprhs(Loomis et al. 2019). Like most other subterranean animals, cave morphs are albino and hypoxia tolerant(O’Gorman et al. 2021; van der Weele and Jeffery 2022).

Apart from a few genetic lesions partly explaining the loss of eye and melanin, not many inactivating mutations in genes have been observed in cavefish(Gore et al. 2018; O’Gorman et al. 2021). The expression divergence of genes through altered regulatory circuitry may contribute to phenotypic divergence. Through this study, we hypothesize that gene regulatory circuitry could have been altered in cave morphs through mutations in the conserved noncoding elements that serve as binding sites of transcription factors. The resulting divergence or loss of gene-expression may have caused the suppression of developmental pathways engaged in regressed phenotypes of cave morphs.

Thousands of conserved non-coding elements (CNEs) have been identified in vertebrate genomes(Elgar 2009). It is established that majority of CNEs serve as the binding sites of DNA-binding proteins, and often colocalize to development related genes(Elgar 2009). Interestingly, teleost had rapid turn-over and divergence of conserved noncoding elements in their evolutionary history(Lee et al. 2011; Ravi and Venkatesh 2018; Iliopoulou et al. 2023). Loss or the divergence of CNEs may link to loss of lineage-specific phenotypes(McLean et al. 2011; Hiller et al. 2012; Marcovitz et al. 2016). We, therefore, tested if the loss of CNEs in cave morphs explains the evolution of regressed and diverged phenotypes of cave morphs.

## Materials and Methods

### Identification of conserved noncoding elements

Genome sequences of Zebrafish (GRCz10), Cave Mexican tetra (GCA_019721115.1), Prianha (GCA_015220715.1) and Surface Mexican tetra (GCA_023375975.1, GCA_000372685.2) were obtained from Ensembl (https://asia.ensembl.org/index.html) and NCBI (https://www.ncbi.nlm.nih.gov/) servers. Pairwise alignment was conducted using Anchorwave software (https://github.com/baoxingsong/AnchorWave), and the resulting ‘maf’ files were converted to ‘axt’ file format(Song et al. 2022). The pairwise ‘axt’ files were processed through the ‘CNE’ function to create a CNE class object using CNEr package (https://rdrr.io/bioc/CNEr/)(Tan et al. 2019). Subsequently, CNEs were identified employing the ‘ceScan’ function with parameters identities=35, window=50 (i.e. at-least 70% sequence identity), and filters for exonic and repeat sequences. CNEs mapped to unaligned regions flanking the anchors were discarded. We extracted the conserved CNEs, cave-lost CNEs, and surface-lost CNEs utilizing ‘intersect’ function of bedtools (https://bedtools.readthedocs.io/en/latest/)(Quinlan and Hall 2010). Proximal genes to these CNEs were identified using bedtools ‘closest’ function, with a distance threshold of 50kb. These genes were subsequently categorized into three groups based on their proximity to CNEs, namely genes near cave-lost CNEs, genes near surface-lost CNEs, and genes near conserved CNEs.

### Analysis of histone modification data

Processed bigwig files for H3K4me3, H3K4me1, H3K27ac, H3K27me3 modifications of 48hpf zebrafish embryo were acquired from DANIO-CODE (https://danio-code.zfin.org/). CNE-centered heatmaps of histone modifications were constructed using deepTools (https://deeptools.readthedocs.io/en/develop/)(Ramírez et al. 2016).

### Hi-C analysis

The KR-normalized .hic files for 48 hpf zebrafish embryos were obtained from GEO (GSE156099). Hi-C matrices at 5 kb resolution were extracted using Juicer software (https://github.com/aidenlab/juicer)(Durand et al. 2016). We extracted the rows corresponding to gene transcription start sites (TSSs) for virtual 4C analysis. We considered only the bins located +/-1 Mb from TSSs. The CNE-gene pairs located in the adjacent bins were not considered. The regions between CNEs and the TSSs were binned relatively into two bins. Similarly, we extended the outer ends of CNEs and TSSs by exactly the same distance as CNE-gene distance and binned those into two bins each. The signals from original 5kb bins within the relative bins were averaged to obtain the comparable values for the relative bins. Heatmaps and boxplots for the virtual 4c signals were drawn using R. We used HiTAD to identify Topologically Associated Domains (TADs) from 48hpf zebrafish embryo Hi-C data(Wang et al. 2017). In-house scripts were used to generate aggregation plots across TAD borders. The SRA files of 48hpf embryo Hi-ChIP (using H3k4me3 antibody) data were retrieved from GSE175706(Franke et al. 2021). We used HiCUP (https://www.bioinformatics.babraham.ac.uk/projects/hicup/) to map the paired reads to reference genome (GRCZ10). The resulting BAM files were fed into FitHiChIP for loop calling (Bhattacharyya et al., 2019).

### Gene Ontology analysis

We used GREAT version 3.0.0 (http://great.stanford.edu/great/public-3.0.0/html/)(McLean et al. 2010) for GO analysis. The enrichment results based on only 1 gene were removed. The terms were sorted based on Benjamini-Hochberg corrected FDR. Expression enrichment analysis was performed using in-situ hybridization data on TopAnat server (https://www.bgee.org/analysis/top-anat).

### Motif analysis

Motif enrichment within cave-lost CNEs, contrasting with surface-lost CNEs, was investigated using ‘motifmatchr’ package (https://www.bioconductor.org/packages/release/bioc/html/motifmatchr.html). We used ‘makeBackground’ function to prepare the background from surface-lost CNEs. ‘motifEnrichment’ function was used for motif enrichment in cave-lost CNEs, taking background of surface-lost CNEs. We performed gene ontology enrichment for the significantly enriched (P<0.05, n=241) transcription factors using ShinyGO (http://bioinformatics.sdstate.edu/go/).

### Gene expression analysis

Developmental gene expression data from Astyanax *mexicanus* was obtained from the supplementary dataset of Stahl & Gross (Stahl & Gross, 2017). DESeq2 (https://bioconductor.org/packages/release/bioc/html/DESeq2.html) was used to compute the log2 fold change in gene expression between cave and surface fish.

The single cell RNA-Seq data for whole zebrafish embryos up to 5dpf were obtained from Farnsworth et al and Farrell et al. (Farrell et al. 2018; Farnsworth et al. 2020). The supplementary table S1 of Farnsworth et al was used to obtain the mean expression value for each cell-type cluster. The UMAP biplots of Farell et al’s data were generated using Daniocell (https://daniocell.nichd.nih.gov/index.html).

### Statistical analysis

To assess statistical significance of differences between distributions, we used non-parametric tests of significance, namely Mann Whitney U tests, on R software. We obtained the false discovery rates (FDRs) though Benjamini-Hochberg method implemented on GREAT, TopAnat, and ShinyGO servers.

## Results and discussion

Availability of high quality genome assemblies for both cave and the surface morphs of A. *mexicanus* enables comparative genomics, facilitating the identification and study of conserved non-coding elements(Warren et al. 2021). By aligning the genomes of D. *rerio*, P. *nattereri*, and both morphs of A. *mexicanus*, we identified 57,484 CNEs conserved across species (Figure 1A-B).. The cave-morph of A. *mexicanus* lost 3,343 CNEs independently, while the surface morph lost 1,056 CNEs (Figure 1B). These CNEs predominantly exhibited enhancer associated chromatin marks, namely H3K27ac and H3K4me1 rather than promoter-associated H3K4me3 or the repression-associated H3K27me3 marks, in zebrafish embryo(Figure 1C), indicating their role as enhancer type distal regulatory elements. CNEs were enriched within the interior of topologically associated domains (TADs), and depleted at the gene-rich TAD-borders (Figure 1D), consistent with their evolutionary synteny and correlation with TAD structure (Harmston et al. 2017). Further analysis using zebrafish embryonic Hi-C data revealed that CNEs were frequently in spatial contact with proximal genes through chromatin loops, supporting their role as distal regulatory elements (Figure 1E). These findings strengthen the assertion that observed CNEs regulate nearby genes.

**Figure 1.**
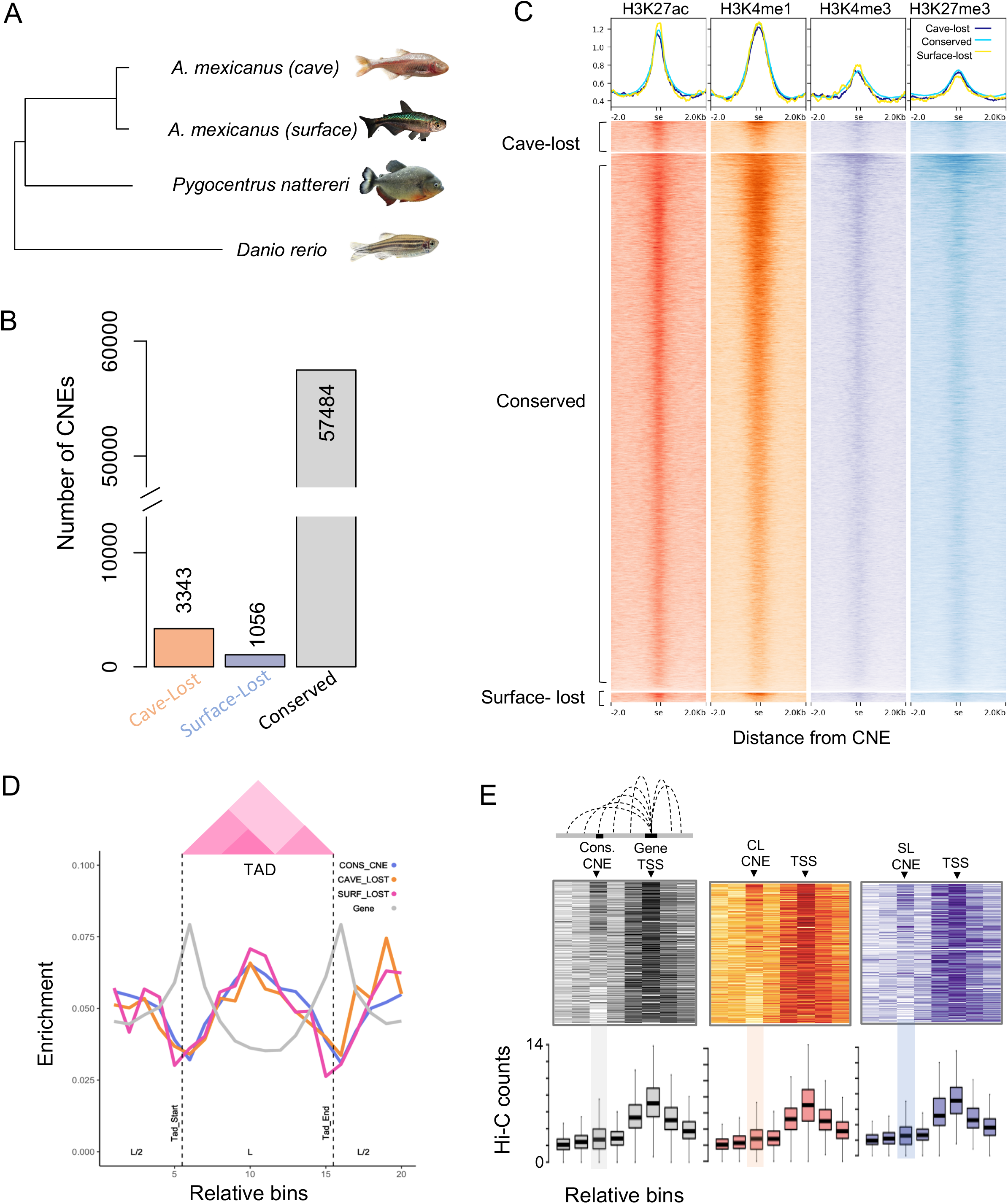
Identification and epigenetic characterization of conserved non-coding elements (CNEs) that were conserved in D. rerio, P. nattereri and surface/cave morphs of A. *mexicanus*. **(A)** Phylogenetic relationships among the four fishes. **(B)** Number of CNEs that were conserved in all four fishes and the lost ones in cave/surface morphs. **(C)** Enrichment of histone modifications H3K4me3, H3K4me1, H3K27ac, and H3K27me3, derived from 48 hpf zebrafish embryos, within and around the conserved, cave-lost and surface-lost CNEs. **(D)** Mean enrichment of conserved, cave-lost, surface-lost CNEs, and the gene TSSs around TADs of 48hpf zebrafish embryos. **(E)** Virtual 4C signals, generated from Hi-C data of 48hpf zebrafish embryos, while taking CNE and the TSS of the proximal gene as reference points.

The genes near the cave-lost CNEs were significantly associated with the eye and nervous system related terms, while the genes near surface-lost CNEs showed enrichment of hypoxia, mesoderm and glycogen metabolism related terms (Figure 2A-B). Coherently, analysis of expression ontology terms derived from in-situ hybridization studies highlighted significant enrichment of nervous system related terms among genes proximal to cave-lost CNEs, while genes near surface-lost CNE did not exhibit enrichment of any term (Figure 2C-D). Elevated expression of genes neighbouring cave-lost CNEs in zebrafish retinal and brain neurons, further confirmed their involvement in eye and nervous system related functions (Figure 3A). We demonstrated the specific expression of *Irx1a* and *Pcdh17*, pivotal genes in teleost retinal development(Cheng et al. 2006)(Chen et al. 2013), in eye and neural cells using single cell transcriptome of zebrafish embryos (Figure 3B). Motif analyses revealed a notable enrichment of binding sites of retina and nervous system related transcription factors in cave-lost CNEs (*vs*. surface-lost CNEs, Figure 3C). Top ranking motifs were those of *Rarg*, Pou3f1, and Foxj3 transcription factors, known to be implicated in eye and nervous system development(Cvekl and Wang 2009)(Fries et al. 2023)(Landgren and Carlsson 2004). Their selectively elevated expression in eye and neural cells, illustrated through single cell transcriptome of zebrafish embryos, further supports our claim that cave-lost CNEs primarily associated with eye and nervous system development (Figure 3D).

**Figure 2.**
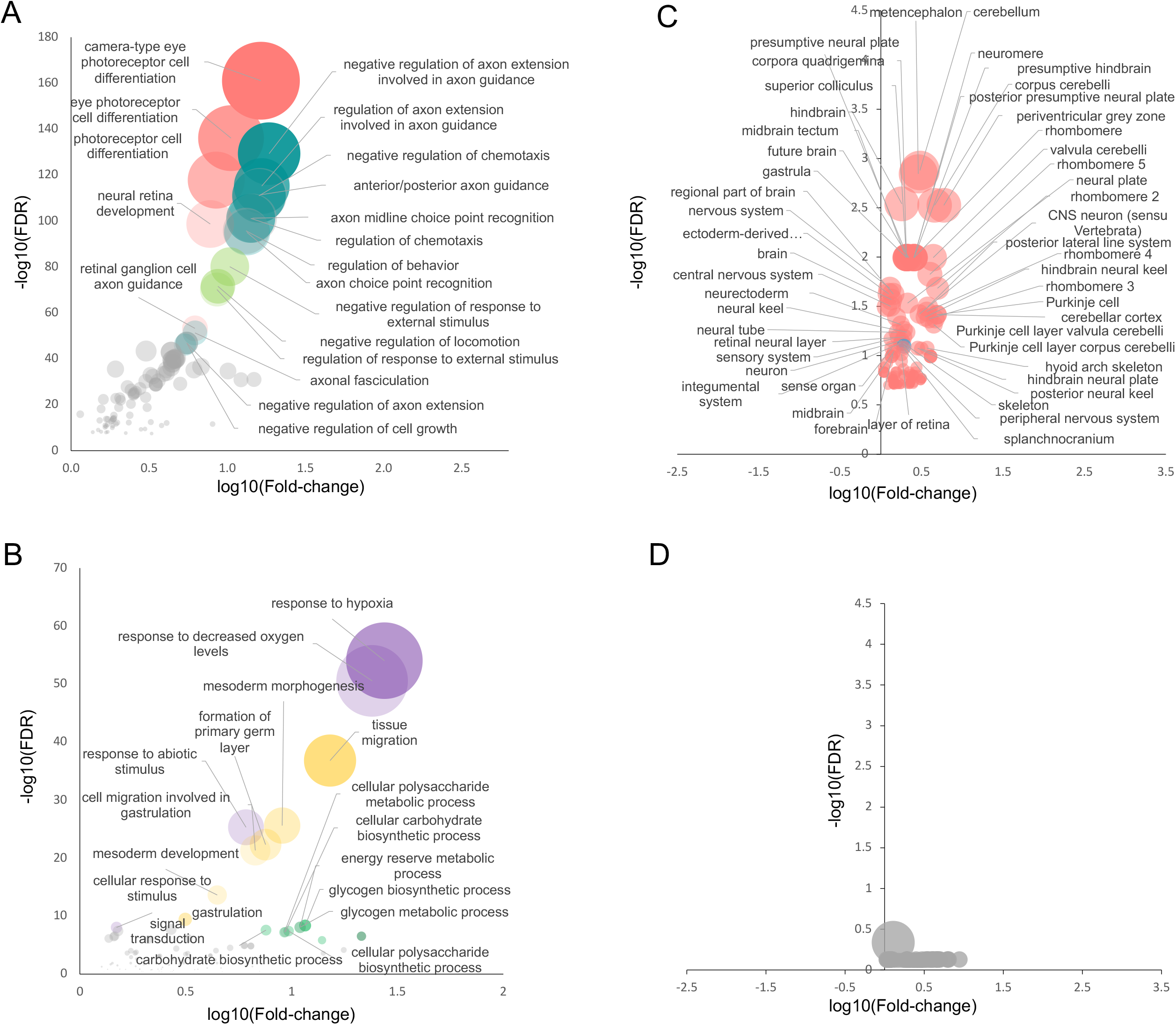
Functional analysis of genes proximal to CNEs. **(A-B)** Enrichment of gene ontology terms among genes proximal to cave-lost and surface-lost CNEs. The genes near conserved CNEs were taken as a background set. FDR is calculated using Benjamini-Hochberg method. Plotted are the volcano plots of FDR as a function of fold enrichment. The diameters of circles are scaled to -log10(FDR). Different functions are arbitrarily coloured. **(C-d)** Gene-expression enrichment analysis using in-situ hybridization data. The FDR of enriched expression ontology terms was plotted as function of fold change over expected background. The genes proximal to surface-lost CNEs did not show enrichment in any particular tissue or cell-type. The P-values were corrected using Benjamini-Hochberg method.

**Figure 3.**
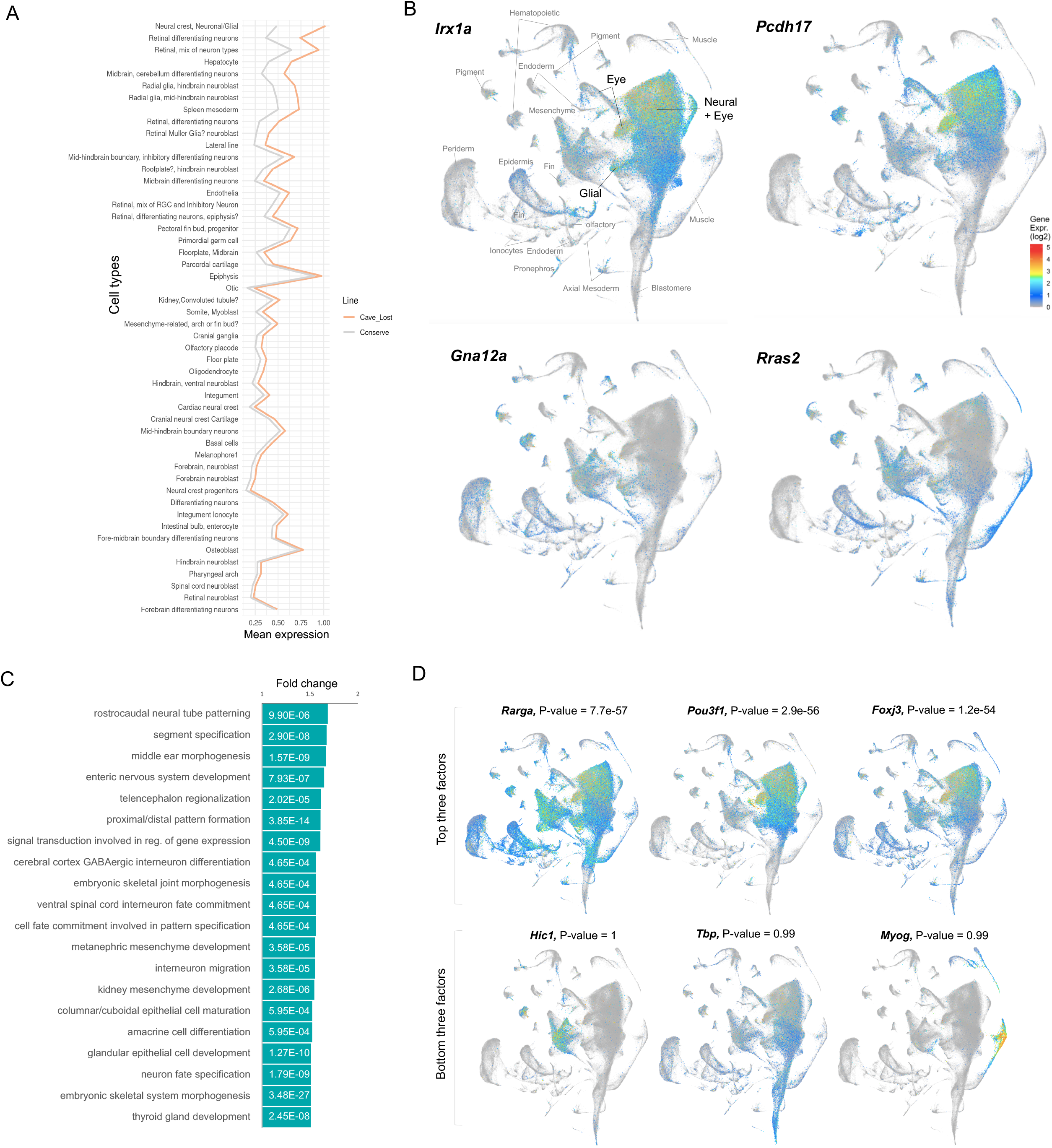
Single cell expression analysis of genes and transcription factors associated with cave-lost CNEs. (**A)** Mean expression levels of nearest genes within 50kb to cave-lost and conserved CNEs across various cell-types of Zebrafish embryos. The cell-types are sorted based on relative difference in mean expression of genes in cave-lost and conserved sets. Only top 50 cell-types are shown. **(B)** Expression of *Irx1a* and *Pcdh17* genes mapped onto UMAP biplot of scRNA-Seq data across various cell-types of zebrafish embryo. The major cell-type clusters are labelled. Expression of example genes (Gna12a, Rras2) from the bottom of the gene ontology enrichment are shown as control in the lower panel **(C)** Enrichment of gene ontology terms for the transcription factors that exhibited enriched binding sites in the cave-lost CNEs. P-values were corrected using Benjamini-Hochberg method. **(D)** Expression of top three and the bottom three transcription factors across single cells of zebrafish embryos.

Since genes near cave-lost CNEs were predominantly associated with regressed phenotypes, we investigated whether the loss of CNEs corresponded to a loss of expression of proximal genes. Analysing developmental time-course gene expression data from cave and surface morphs, we observed progressive downregulation of genes near cave-lost CNEs during embryonic development compared to those near conserved and surface-lost CNEs (Figure 4A-B). This downregulation coincided with the post-segmentation eye growth and brain patterning stages, typically around 36 hours post-fertilization, when eye growth is known to be arrested in cave morphs (Jeffery and Martasian 1998; Jeffery et al. 2003; Alunni et al. 2007). These analyses suggest that the loss of CNEs likely modulated developmental pathways related to the eye and nervous system in cave morphs, potentially contributing to the regression of associated phenotypes. Indeed, previous studies have shown that loss of CNEs is associated with loss of certain phenotypes in mammals (Hiller et al. 2012; Marcovitz et al. 2016).

**Figure 4.**
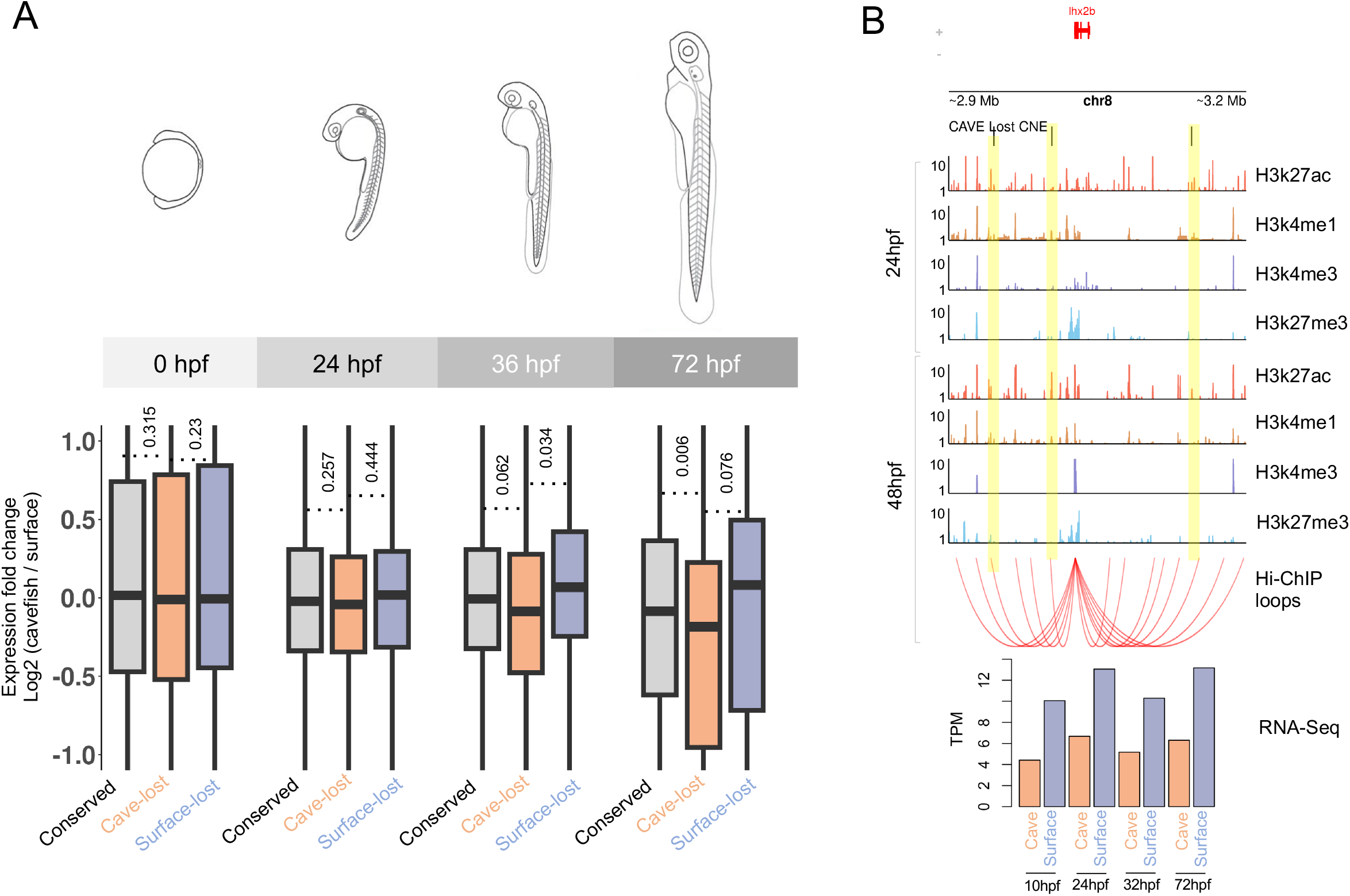
Developmental expression of genes proximal to CNEs. **(A)** Expression of genes near cave-lost, surface-lost, and conserved CNEs across four developmental stages of Zebrafish embryo. P-values for the lower expression in the cave-lost set in the given comparisons were calculated using Mann-Whitney U tests. **(B)** An example of cave-lost CNEs, near gene *Lhx2b*, associated histone modifications, chromatin loops, and gene expression in Zebrafish embryos.

Interpretating the gene ontology terms for the surface-lost CNEs is convoluted. These CNEs were flanked by genes related to hypoxia, mesoderm formation, and glycogen metabolism terms. Notably: i) Cave morphs are more tolerant to hypoxia than surface morphs(van der Weele and Jeffery 2022); ii) Cave morph embryos display earlier convergences, extension and internalization of mesodermal cells(Torres-Paz et al. 2019); iii) Cave morphs have enhanced glycogen metabolism than surface morphs(Olsen et al. 2023). Interestingly, these three terms-hypoxia, mesoderm and glycogen metabolism - are inter-connected, as hypoxia can stimulates both the glycogen metabolism and the blood cell formation. These results may imply that the surface morph had lost the CNE near genes that were associated with the constructive traits of cave-morphs. Although D. *rerio*, P. *nattereri* are not subterranean, the surface-lost CNEs were present in these species. This possibly implies that some surface-lost CNEs were evolutionarily constrained in other fishes included cave morphs, but not in surface morphs. It is worth noticing that the zebrafish shows one of the lowest P_crit_ values, among 195 fishes analysed in a study, marking significant hypoxia tolerance(Rogers et al. 2016). While such data could not be retrieved for surface morph, studies suggest that haemoglobin concentration (∼10.5 g/dl) and the haematocrit level (∼35%) of cave morphs matches with that of zebrafish (∼10 g/dl and 31%), while surface morphs have significantly lower values (8.25 g/dl and 28%). We, therefore, hypothesize that the surface morph of A. *mexicanus* may have less tolerance to hypoxia when compared to zebrafish. Cave morph, on the other hand, retained the CNEs that governs the hypoxia tolerance owing to its adaptation to hypoxic environment.

In summary, our observations highlighted the genetic underpinning of regressed phenotypes in cave-morphs of Astyanax *mexicanus*. The loss of conserved noncoding elements associated with the suppression of eye and neuronal pathways during embryonic development of cave-morph, likely shaping the evolutionary regression of allied phenotypes in cave-morphs. These observations also provide valuable insights for designing genetic experiments, such CNE knock-outs in surface morphs, to further unravel evolutionary pathways driving cave adaptations.

## Acknowledgement

The authors duly acknowledge the financial support from Department of Biotechnology, India and IISER-Mohali.

## Funding

This work was funded by Department of Biotechnology, India (BT/PR40149/BTIS/137/36/2022, BT/PR40198/BTIS/137/56/2023).

## Data availability

All associated datasets are available as supplementary material.

## Conflict of interests

Authors declare no competing interests.

## Supplementary Datasets

**Dataset S1**. Chromosomal coordinates for the Conserved, cave-lost and surface-lost CNEs in the zebrafish genome assembly GRCz9. Gene-ontology terms can be reproduced by submitting these coordinates on GREAT server.

**Dataset S2**. Data for the histone modifications around conserved, cave-lost, and surface-lost CNEs.

**Dataset S3**. Virtual 4c data for the CNEs and the cognate gene TSSs for the conserved, cave-lost, and surface-lost sets.

**Dataset S4**. Enrichment of gene ontology terms in the genes proximal to conserved, cave-lost, surface-lost CNEs.

**Dataset S5**. Cell-type specific and developmental expression data for genes proximal to CNEs.

**Dataset S6**. Motifs enriched in cave-lost CNEs, contrasted with surface-lost CNEs.

## Notes

### Competing Interest Statement

The authors have declared no competing interest.

